# The Cupin Protein, Dehydratase Pac13 is a Homodimer

**DOI:** 10.1101/600247

**Authors:** Rundong Zhang

## Abstract

Cupin proteins share a double-stranded β-helix fold, form one of the largest superfamilies and possess remarkable functional diversity. They usually form homooligomeric states. Michailidou et al. recently reported that a cupin protein Pac13, which is a dehydratase mediating the formation of the 3’-deoxy nucleoside of pacidamycins, is an unusual, small monomer. However, a careful analysis of the biophysical and structural data provided by the authors clearly indicates that Pac13 is a homodimer.

---

Cupin-like proteins share a double-stranded β-helix fold, form one of the largest superfamilies and possess remarkable functional diversity^[1]^. Many of them are involved in metabolism of various antibiotics and compounds with biological activities, and therefore structural characterizations of them, including their oligomeric states, are important. Cupin proteins usually form homo-oligomeric states^[1]^. However, Michailidou et al. recently reported that a cupin protein Pac13, which is a dehydratase mediating the formation of the 3’-deoxy nucleoside of pacidamycins, is a small monomer^[2]^. They claimed that Pac13 is an unusual, small and monomeric dehydratase. Their statement of Pac13 as a monomer is one of their major conclusions as it is emphasized multiple times in the title, abstract, main text and supporting information^[2]^. However, a careful analysis of the biophysical and structural data provided by the authors clearly indicates that Pac13 is a homodimer.

The full-length Pac13 contains 121 residues with Molecular Weight (MW) of 13815 Da (UniProtKB – E2EKP5). From the “protein production and purification” section in the supporting information^[2]^, it can be known that the studied construct contained the full-length Pac13 with a His_8_ tag followed by a TEV cleavage site fused at its N-terminus. During the purification, the His_8_ tag was removed by incubation with TEV protease before size exclusion chromatography (SEC). The tag-removed Pac13 protein on the SDS-PAGE was at about 15 kDa (Figure S1 in the original paper^[2]^) and the calculated MW of wildtype Pac13 is 14096.67 Da (Table S1 in the original paper^[2]^). The protein standards from SEC column HiLoad 16/600 Superdex 75 prep grade column gave rise to the equation: y=4111.2e^−0.071x^, in which y is MW in kDa, and x is the elution volume in ml (Figure S2 in the original paper^[2]^). The major peak of Pac13 from the same column was about 68.8 ml (with the confidence within the range of 68 to 70 ml, Figure S2 in the original paper^[2]^), then the calculated MW is 31.08 kDa and about 2.2 times of a monomer’s MW (with the confidence within the range of 28.54 to 32.90 kDa, 2.0-2.3 times of a monomer’s MW). Therefore, the studied Pac13 protein in solution should be a homodimer.

In addition, from the crystal data (PDB codes: 5NJO and 5NJN) determined by the paper^[2]^, it should be deduced that Pac13 is a homodimer. Although Pac13 crystallized in space group P3_2_21 with one molecule in the asymmetric unit, it is clearly seen that each molecule in the unit cell has a closely contacted neighbor and they form a pair in C2 symmetry (Figure 1). The analysis by PISA (Proteins, Interfaces, Structures and Assemblies)^[3]^ (http://www.ebi.ac.uk/msd-srv/prot_int/) also indicates that Pac13 is a homodimer (Supplemental Figure 1). Taking the coordinate of PDB code 5NJO as an example, the interface area for each subunit is 1392.3 Å^2^ and the total buried area is 2784.5 Å^2^. There are 17 H-bonds, 2 salt bridges and a number of hydrophobic interactions between the interfaces. The predicted free energy gain upon formation of the homodimer (ΔG^int^) is −17 kcal/mol and the free energy of assembly dissociation (ΔG^diss^) is 13 kcal/mol. Using the equation, ΔG^diss^= −RTlnK_d_, and the temperature of 25°C, the dissociation constant K_d_ can be calculated to be 2.9×10^−10^ M (the association constant K_a_ should be 3.0×10^12^ M^−1^, using the above ΔG^int^ value), which is within the tightest noncovalent interactions for biomolecules. The Complexation Significance Score (CSS), which indicates how significant for assembly formation the interface is, is 1, the highest value, and the PISA server predicts that the dimeric assembly is stable.

**Figure 1.**
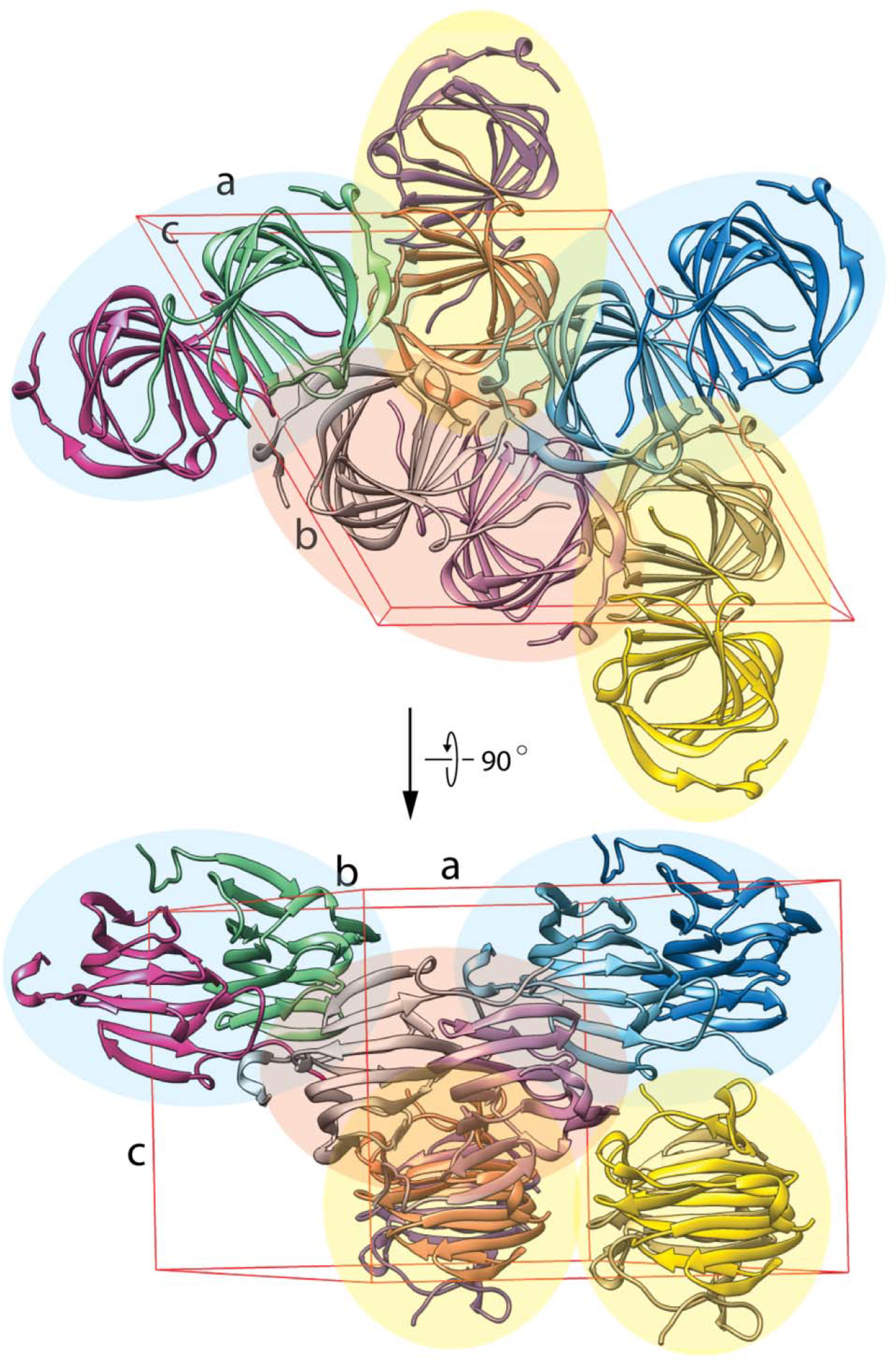
Pac13 is a homodimer in crystal. The unit cell of space group P3_2_21 (PDB code: 5NJO) is shown in red. Each of the six monomers inside the unit cell forms a tightly contacted pair with its neighbor in C2 symmetry (in elliptic shadows colored in blue, yellow and red). Each monomer is show in cartoon and colored differently.

In the homodimer of Pac13, the interactions between each subunit are mediated mainly by the anti-paralleled 6-stranded β sheet (β1-β2-β3-β10-β5-β8), termed here as the inner sheet (Figure 2). The anti-paralleled 5-stranded β sheet (β7-β6-β9-β4-β11) is located outside of the dimer and termed as the outer sheet. In addition to the cupin-fold core consisting of 11 β-strands, which was reported in the original paper, the N-terminus of each subunit contains an additional β strand, termed β0 (residues 4-6), which interacts with β8 of the second subunit in anti-parallel, and extends the inner sheet (Figure 2). Moreover, each β0 also interacts with β7 and β8 from the other subunit with hydrophobic interactions (Figure 2). In addition, the interactions between the inner sheets are mostly mediated by the four β-strands, β3-β10-β5-β8, and are largely mediated by hydrophobic interactions (Figure 2). V33 from β3, V96 from β10, V58 from β5, and L78 from β8 form the major hydrophobic surfaces on the inner sheets for packing into a dimer. There are also a few hydrogen-bonding interactions interspersed among the hydrophobic interface, for example, between the side chains of the two T98 and between the side chains of Y56 and R35 (not shown). The detailed analysis of the interactions also indicates that these inter-subunit interactions are specific and extensive, and therefore, Pac13 should form a stable homodimer.

**Figure 2.**
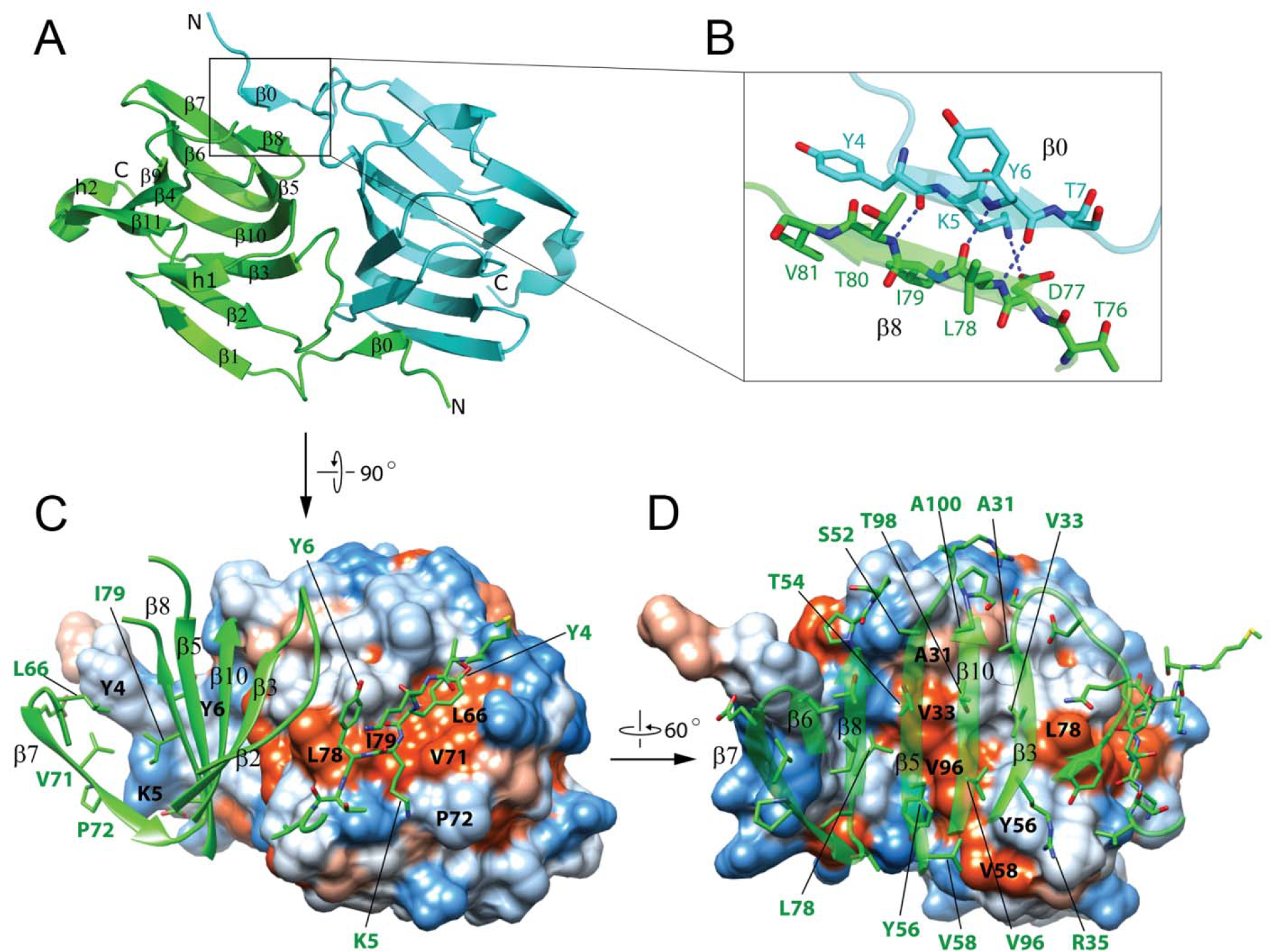
The interactions at the interfaces of Pac13 homodimer are specific and extensive. (A) One Pac13 homodimer. The two monomers are shown in cartoon and colored in green and cyan respectively. The β strands (β1-β11) and helices (h1-h2) are labelled. N and C indicate the N- and C-termini respectively. (B) The detailed view of the anti-parallel interaction between strands β0 and β8 (from different subunits). H-bonds are shown in blue dashed lines. (C) Each β0 (Y4, K5 and Y6) also interacts with β7 (L66, V71 and P72) and β8 (L78 and I79) from the other subunit with hydrophobic interactions. (D) The interactions between subunits are mediated mostly by the four β-strands, β3-β10-β5-β8, of the inner sheets, and are mainly hydrophobic interactions. In (C) and (D), one subunit is shown in surface with each residue’s hydrophobicity mapped on it in different colors (colors from orange to blue indicate hydrophobicity from high to low). For clarity, the segments without inter-subunit contact in the green subunit are not shown.

The authors also claimed that “Pac13 remains the smallest cupin with a characterized enzymatic activity” (page 26 in the supporting information in the original paper^[2]^). If Pac13 were a monomer, this statement might be right. However, because Pac13 is a homodimer, this statement is invalid. There is another cupin-fold enzyme reported earlier than the Pac13 paper, Kdgf (PDB code: 5FPX)^[4]^, which is a homodimer of 114-residue protein with MW of 12821 Da and smaller than Pac13.

Interestingly, there are many cupin-fold proteins forming homodimers with the inter-subunit interactions similar to those of the Pac13 homodimer. Take a couple of them mentioned in the Pac13 paper as examples: the lyase KdgF (PDB code: 5FPX)^[4]^, which is the closest homologue with characterized function to Pac13 using the Dali server, and the ectoine synthase (EtcC) (PDB code: 5BXX)^[5]^, which is reported as the only other cupin protein shown function as dehydratase, are both homodimer, with similar interaction patterns between their subunits to those of the Pac13 homodimer (Figure 3. compare with Figure 2). They all form homodimers with close contacts by the inner sheets and each β0 in one subunit interacts in anti-parallel with β8 of the inner sheet of the other subunit. Also interestingly, both these cupin-fold enzymes, although not always crystalizing in the same space groups, do crystalize in the same space group of P3_2_21 as does Pac13. In the case of KdgF (from *Yersinia enterocolitica*), it forms space group P3_2_21 in PDB code of 5FPZ, in addition to space group P2_1_2_1_2_1_ (PDB code: 5FPX)^[4]^. EtcC (from *Sphingopyxis alaskensis*) forms space group P3_2_21 in PDB code of 5BY5, in addition to space group C2 (PDB code: 5BXX)^[5]^. Our recent crystallographic study of another cupin-fold enzyme also crystallized in space group P3_2_21, and the enzyme is also a homodimer with the above-mentioned inter-subunit interactions (data not shown, the manuscript in preparation).

**Figure 3.**
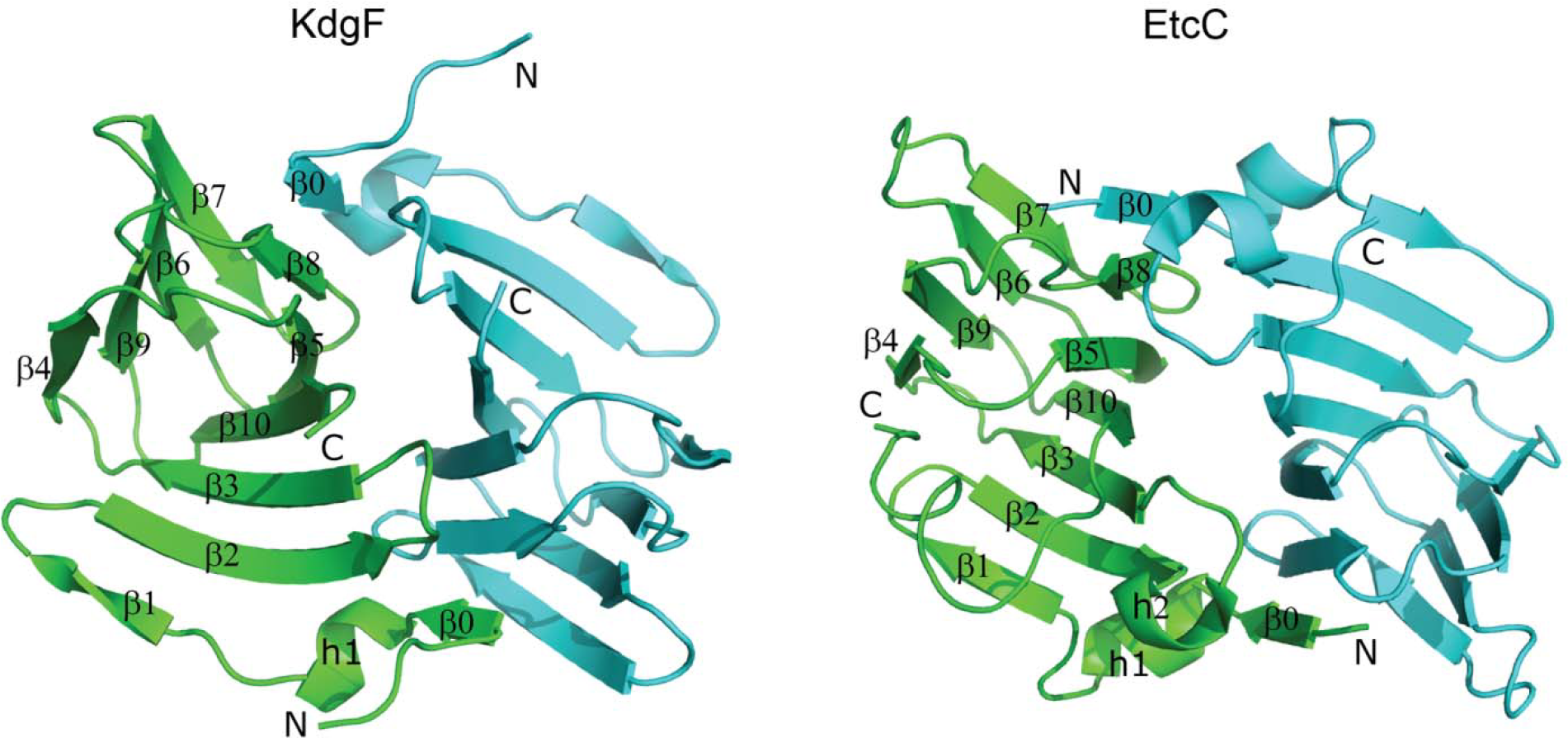
Other cupin proteins form similar homodimers. The lyase KdgF (PDB code: 5FPX) and the ectoine synthase EtcC (PDB code: 5BXX) form homodimers similar to Pac13.

In conclusion, a careful analysis of the Pac13 data from the original paper concludes that Pac13 is a homodimer both in solution and crystal and it is also not the smallest cupin protein. The statement of the authors that Pac13 is an unusual monomeric dehydratase^[2]^ is invalid.

## ACKNOWLEDGEMENTS

This work was supported by National Key R&D programs (No. 2017YFA0505800 and 2017YFA0504300) and National Natural Science Foundation of China (No.31570720).

