## Supplemental Figure 1 for "The Cupin Protein, Dehydratase Pac13 is a Homodimer"

Rundong Zhang*

**A**

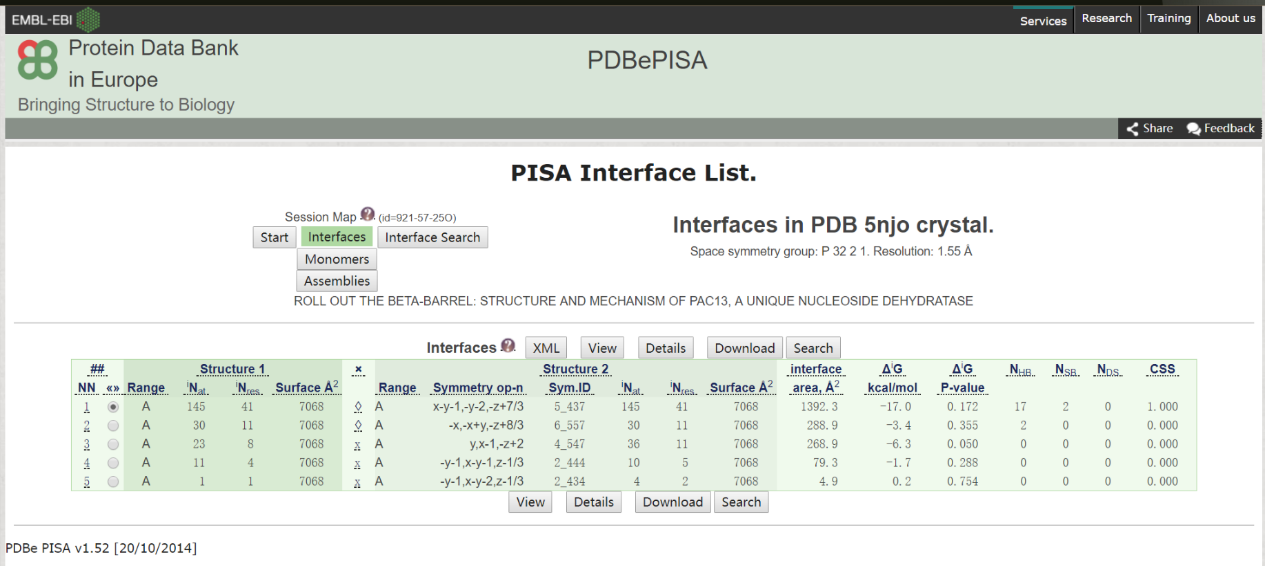

**B**

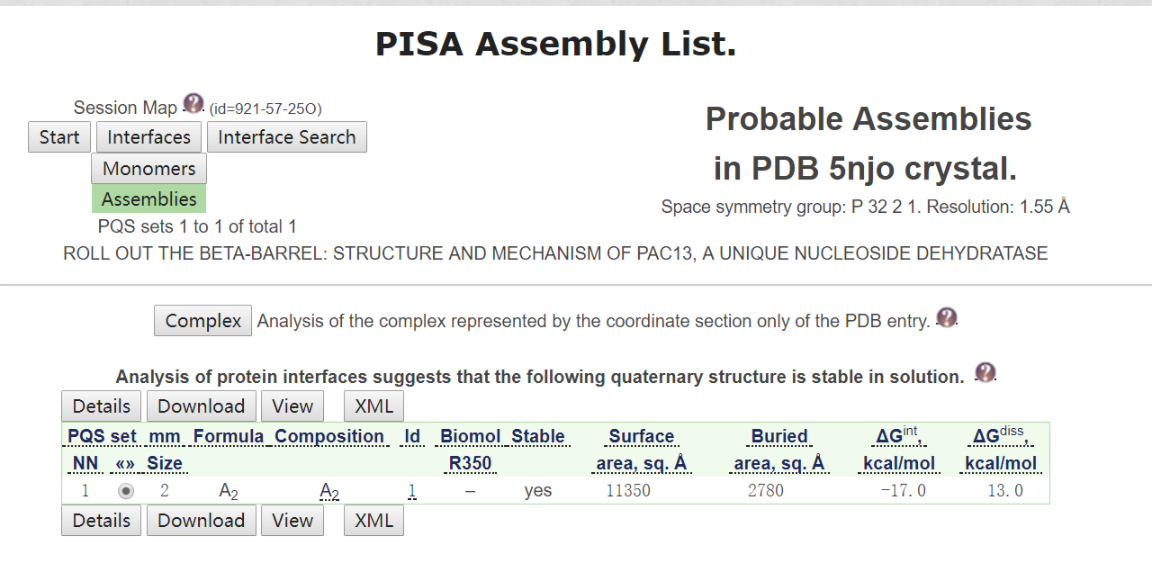

**Supplemental Figure 1. The PISA analysis of Pac13 coordinate of PDB code 5NJO.** (A) Interface List. (B) Assembly List.
